# Perception of first and second pain during offset analgesia

**DOI:** 10.1101/2025.06.30.662291

**Authors:** Jakob Poehlmann, Benita von Lemm, Luisa Luebke, Waclaw M. Adamczyk, Kerstin Luedtke, Tibor M. Szikszay

## Abstract

**Introduction:** Offset analgesia (OA) is defined as a disproportionate reduction in pain perception following a small decrease in noxious stimulation. However, the mechanisms underlying this phenomenon remain unclear, with ongoing debate on peripheral versus central contributions.

**Objectives:** This experimental study aimed to differentiate first and second pain perception during the OA paradigm, thereby assessing fiber-specific influences on OA.

**Methods:** Thirty-two healthy participants were asked to distinguish a double pain sensation (first and second pain), to assess pain quality descriptors related to A-δ and C-fibers, and to indicate response times to brief noxious heat stimuli. This procedure was repeated while implementing heat pulses in an OA paradigm.

**Results:** No significant differences were found between offset and constant trials in the reported double pain sensation or the fiber specific pain descriptors (p > 0.05). Nevertheless, significant differences in response times were observed depending on the type of trial and the timing of the stimulus. Response time to noxious stimuli was delayed after prolonged stimulation in both offset and constant trials (p < 0.05).

**Conclusions:** The findings suggest that A-δ and C-fiber response characteristics were unaffected during the OA paradigm; however, higher stimulation intensities or prolonged pain induce a notable response delay. This indicates a negligible role of specific peripheral nerve fibers in OA, emphasizing the predominance of central mechanisms, particularly those related to attention and cognitive resources, which merit further investigation.

## 1. Introduction

Offset analgesia (OA) is characterized by a disproportionate decrease in reported pain intensity following a minimal reduction of noxious stimulation (Grill & Coghill, 2002). To understand the mechanisms underlying this phenomenon, extensive research using brain imaging has demonstrated the central processing of OA in brain regions associated with descending pain modulation (Derbyshire & Osborn, 2009; Ito et al., 2024; Li et al., 2022; Nahman-Averbuch et al., 2014; Sprenger et al., 2018; Yelle et al., 2009; Zhang et al., 2018). However, less attention has been directed to peripheral influences on OA, although, demyelination of fast-conducting A-δ fibers or neuropathy of small fibers may explain impairments of OA in the elderly and patients with neuropathic pain or fibromyalgia (Hermans et al., 2016; Szikszay et al., 2019). Indeed, there is additional evidence indicating the involvement of peripheral processing in OA. For instance, OA has been observed in hairy skin areas, unlike hairless regions, which may lack low-threshold A-delta (A-δ) afferents (Asplund et al., 2021; Naugle et al., 2013; Szikszay et al., 2022). Furthermore, OA can be elicited by both noxious heat and innocuous cold thermal stimuli, but not by innocuous warm temperatures, suggesting thermospecific activation of A-fiber afferents, rather than C-fiber afferents (Luebke, Von Selle, et al., 2024).

A distinction is made between different types of nociceptors that are sensitive to a different range of stimuli (Dubin & Patapoutian, 2010; Smith & Lewin, 2009). Rapid-onset pain is predominantly mediated by A-δ-mechano-heat-sensitive nociceptors (Gold & Gebhart, 2010). In contrast, unmyelinated small diameter C-fibers respond to thermal, mechanical, and chemical stimuli in a polymodal delayed manner (Sneddon, 2018). The activation of functionally distinct cutaneous nociceptor populations resulted in a variety of pain qualities depending on modality, location, and temporal characteristics (Besson & Chaouch, 1987; Dubin & Patapoutian, 2010; Gold & Gebhart, 2010). A first, well localized, sharp, and stabbing pain (Campbell & LaMotte, 1983; Price, Donald D.; Dubner, 1977) is followed by a second pain that is usually diffuse, throbbing, and burning (Besson & Chaouch, 1987; Price, Donald D.; Dubner, 1977). Additionally, these characteristics are supported by varying response times to fiber specific stimulation as shown previously, where the myelinated A-δ fibers result in lower response times compared to the unmyelinated small diameter C-fibers(Yarnitsky & Ochoa, 1991).

Based on these previously demonstrated properties of different cutaneous nociceptors, the aim of this study is to investigate the peripheral aspects of OA by characterizing first and second pain during the OA paradigm. First, single brief noxious heat pulses will be presented to illustrate and familiarize participants with the distinction between first and second pain. For this the qualitative properties and response times to increasing stimulus intensity will be assessed. Building upon this, the highest heat pulse will be integrated into an OA paradigm, which will follow the same procedure of assessing qualitative as well as quantitative properties of pain perception. Based on a potential inhibition of A- δ fibers during OA (Greffrath et al., 2007; Peng et al., 2003; Schwarz et al., 2000), it was hypothesized that first and second pain are not distinguishable during OA and that the response times after the temperature offset are accordingly delayed.

## 2. Methods

### 2.1. Study procedure

This randomized, within-subjects study was conducted to investigate first and second pain characteristics of brief heat pulses in the context of an OA paradigm. The study was approved by the Ethics Committee of the University of Lübeck (2023-633) and conducted in accordance with the Declaration of Helsinki. The methodology was preregistered on the Open Science Framework (OSF; https://osf.io/st5gd). The assessment was divided into four segments: distinction of first and second pain (1); a rating of single heat pulses regarding pain quality, pain intensity and response time (RT) (2); continuous pain rating of an OA paradigm (3) and OA including heat pulses (4). An overview of the study procedure is provided in **Figure 1a**.

**Fig 1.**
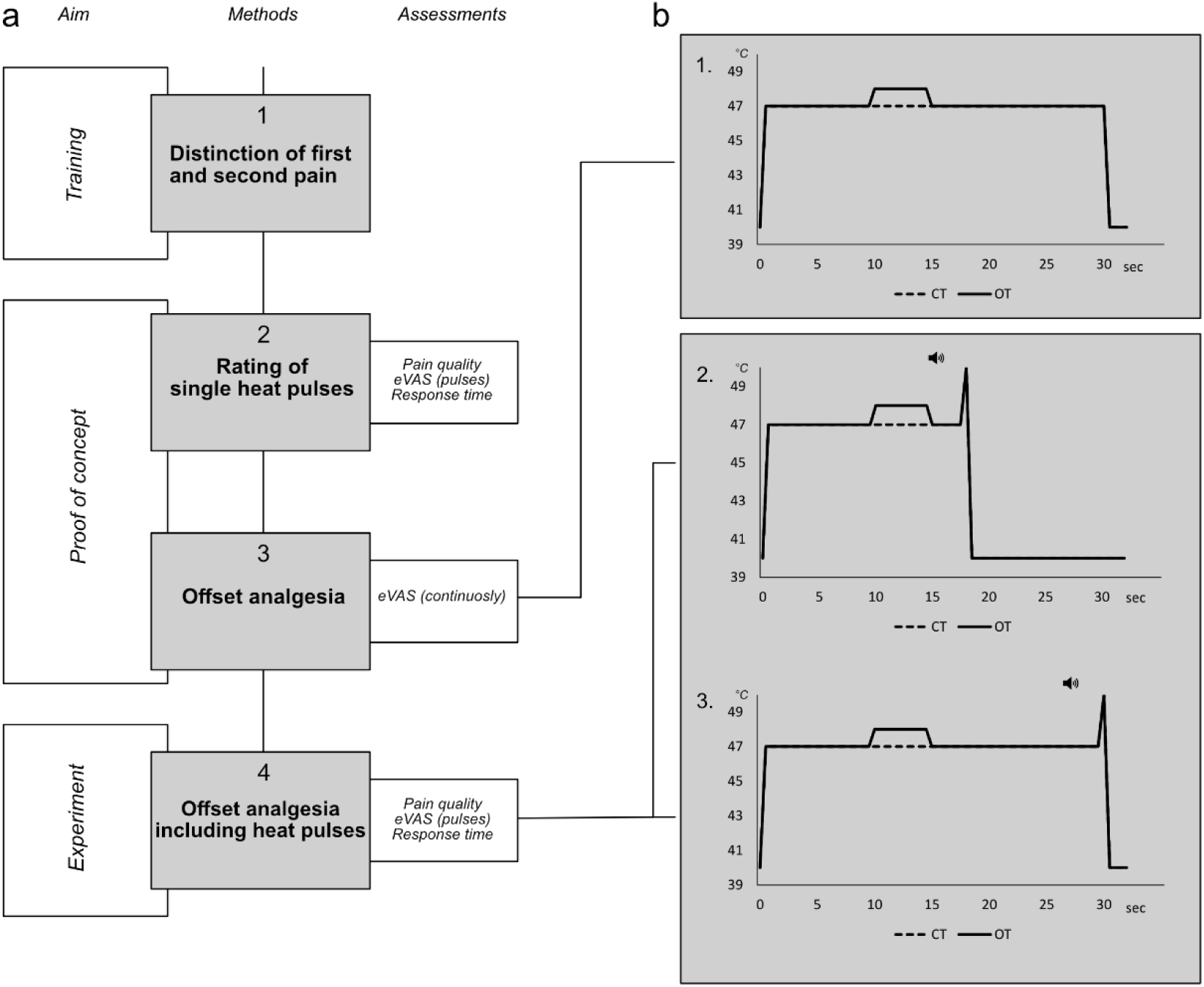
Schematic representation of the study (a) and the experimental paradigm (b) **a)** The study was structured into four stages: (1) Distinction of first and second pain, (2) rating of single heat pulses based on pain quality (double sensations and pain descriptors), pain intensity with an electronic Visual Analogue Scale (eVAS), and reaction time (RT), (3) testing the offset analgesia paradigm using continuous pain rating during offset trials (OT) and constant trials (CT), and (4) offset analgesia including heat pulses measuring pain quality, pain intensity (eVAS) and RT. **b)** Offset trials (OT) and constant trials (CT) without heat pulses (1.), OT and CT with a heat pulse (150 milliseconds, 100°C/second) two (2.) and fifteen (3.) seconds after temperature offset. Upcoming pulses were cued with an acoustic signal.

### 2.2. Participants

Participants (aged between 18 and 45) were included if they subjectively reported being healthy and pain-free on the day of the procedure. Exclusion criteria were chronic pain (> 3 months) within the last two years or diagnosed systemic, neurological, cardiovascular or psychiatric diseases. All participants were asked not to take any pain medication, consume alcohol or undertake any strenuous physical activity 24 hours before the participating in the study. To characterize the study sample, age, sex at birth, body mass (kg), stature (cm), handedness and fear of heat pain on a numerical rating scale from 0 (no fear) to 100 (highest possible fear) were recorded for each participant. To measure two major components of attention (attention focusing and attention shifting) the Attention Control Scale (ACS) was used (Derryberry & Reed, 2002). In addition, the Pain Vigilance and Awareness Questionnaire (PVAQ) was used to measure attention to pain and assesses awareness, consciousness, vigilance, and observation of pain (McCracken, 1997) and the Pain Sensitivity Questionnaire (PSQ) was used to quantify pain intensity ratings of daily life situation (Ruscheweyh et al., 2009).

### 2.3. Equipment

All heat stimuli were applied with a thermal contact stimulator (TCS; André Dufour, University of Strasbourg, France). The probe of the TCS has a total stimulation zone of 8.88 cm^2^ (five equal stimulation zones, each 0.74 × 2.4cm) and was applied to the forearm. The probe weighs 440g and the TCS has a temperature range of 0°C to 60°C, adjustable at 0.1°C intervals. The maximum temperature rise and fall rate is 100°C/second. The reaction time was determined by using a response button with the dominant hand connected to the TCS. Measurement of pain intensity was conducted with a Python-based electronic visual analog scale (eVAS) via a computer, ranging from 0 “no pain” to 100 “worst heat pain imaginable”. Participants were asked to rate the intensity of pain with the dominant hand.

### 2.4. Distinction training for first and second pain

Each participant was instructed to follow a Microsoft PowerPoint presentation (Microsoft Corp., Remond, WA, USA) that described the study procedure and the phenomenon of first and second pain with written and visual information (see **Appendix 1**). The familiarization consisted of eight ascending heat pulses (150 ms, 100°C/s) with intensities from 48°C to 55°C, starting from a baseline temperature of 40°C. The probe was manually placed at three previously marked distinct areas on the dominant ventral forearm. Participants were asked to recapitulate first and second pain described in the presentation while being provided a chart with specific pain descriptors for first and second pain. A previous literature review identified pain descriptors and characterizations of heat-induced primary and secondary pain (see **Appendix 2**).

### 2.5. Rating of single heat pulses

Eight heat pulses (150 milliseconds, 100°/second, baseline temperature 40°C) with temperatures of 49°C, 51°C, 53°C or 55°C (each temperature twice) were applied in a pseudorandomized order. Participants were asked whether they felt a first and second pain (double sensation: yes) or just a single pain (double sensation: no) and to which pain description the pain sensations could be assigned. Participants were allowed to choose once pain descriptor per first or second pain. If they were not able to perceive a double sensation, only one descriptor was chosen. Additionally, they were asked to rate their pain intensity on a numerical rating scale (NRS) of 0-100. After a two-minute break, the participants received eight further identical heat stimuli. As soon as a first pain was perceived, participants pressed a response button (response time).

### 2.6. Offset analgesia

An OA paradigm with three successive periods (T1-T2-T3), consisting of an offset trial (OT) and a constant trial (CT), was performed (Szikszay et al., 2019). CTs had a duration of 30 seconds continuous heat stimulation with an intensity of 46.0 °C. OTs consisted of an intensity of 46.0°C applied for 10s (T1), followed by five seconds of 47.5 °C (T2) and ended with a temperature of 46.0°C for 15 seconds (T3). Rise and fall rates were kept constant (100°C/s). Each trial was performed twice and followed by a break of 2 minutes. The order of trials was pseudorandomized in a counterbalanced manner. Participants were asked to continuously rate the perceived pain intensity. **Figure 1b** indicates a schematic representation of the individual trials.

### 2.7. Offset analgesia including heat pulses

In the following, a heat pulse was integrated into the previously described OA paradigm. This pulse (150 ms, 100°C/s) was applied during T3, either after two seconds or at the end (after 15 seconds) in both the OTs (OT_2s_ or OT_15s_) and CTs (CT_2s_ or CT_15s_). These different trials were chosen to respect for the duration of a possible offset effect (Yelle et al., 2008) (OTs) and control for the duration of nociceptive input with a tonic heat trial (CTs). Each of these trials was performed twice (a total of 8 trials in a pseudorandomized order). In each trial, an acoustic cue was provided one second before the actual pulse. Participants were asked to focus on the integrated heat pulse in the T3 interval of the OA paradigm after an acoustic signal and to rate the pain quality (pain descriptors, double sensation) immediately after the pulse, as previously described. Similarly to the single heat pulses, participants were additionally asked to rate their perceived pain from the heat pulse on a NRS scale of 0-100. In a further four identical trials, the participants were asked to indicate the response time regarding the first pain. See **Figure 1b** for a schematic representation.

### 2.8. Statistical analysis

Sample size calculation was based on the effect size from an earlier OA experiment (Szikszay et al., 2019). With an effect size of d_z_ = 0.76 (α = 5%, β = 95%) (Grill & Coghill, 2002), an estimated group size of n = 25 participants for paired comparisons (two-sided t-test) was required (G*Power, University of Düsseldorf) (Faul et al., 2007). To ensure that the sample size was sufficient to achieve a sampling distribution considered approximately normal, this sample size was increased to n = 32 (Anderson, 2010).

Statistical analyses were performed using R Studio (RStudio version 2024.04.24 with R version 4.4.0, R Foundation for Statistical Computing, Vienna, Austria) (R Core Team, 2024). Mixed-effects models were implemented using the lmer function from the lme4 package (Bates et al., 2015). Parametric data is presented in means (x̄) with standard deviations (SD) and nonparametric data in median (M) with min-max ranges (R) or absolute and relative frequencies. The average eVAS ratings were extracted for each time interval (T1, T2 and T3). The first 5 seconds of T1 and T3 were not considered in our analysis due to the ratings still showing an increasing (T1) or decreasing (T3) ramp. The difference between CT and OT at T3 was considered the OA response, which was analysed by a two-tailed dependent student’s t-test. Regarding OA including heat pulses, the number of double sensations (between 0 and 2) and the amount of fiber-associated pain descriptions (for the first pain) were compared between OT and the CT using the Wilcoxon test. To investigate RT, a generalized linear model (GLM) with the modelled factors „trial“ (OT, CT) and „time“ (2sec, 15sec), was used. If there were significant findings, Bonferroni-corrected t-tests were performed. A similar approach was chosen for the perceived pain intensity (NRS) of the first and second pain. The level of significance was set at p < 0.05.

## 3. Results

### 3.1. Participant characteristics

A total of thirty-two healthy volunteers were recruited in this study. One participant was excluded from the analysis specific to RT, due to showing RTs greater than two standard deviations above the mean in all trials except for one. All participants successfully absolved the distinction training for first and second pain, tolerated the stimulation intensities and showed no signs of adverse events. The demographic characteristics are shown in **Table 1**.

**Table. 1.** Participant characteristics.

| Characteristics n = 32 |  |  |
| --- | --- | --- |
| **Age** | Years, $\bar{x}$ (SD) | 26.7 (4.3) |
| **Height** | Cm, $\bar{x}$ (SD) | 174.6 (10.0) |
| **Weight** | Kg, $\bar{x}$ (SD) | 73.1 (15.0) |
| **Female** | N (%) | 17 (53.1) |
| **Right-handed** | N (%) | 30 (93.8) |
| **Fear of heat pain** | eVAS, $\bar{x}$ (SD) | 27.7(21.8) |
| **Questionnaire** |  |  |
| **ATQ** | M (R) | 46 (42-59) |
| **PVAQ** |  | 35.5 (14-59) |
| **PSQ** |  | 43.5 (25-87) |

*ATQ, Attention control scale; M, median; PSQ, Pain sensitivity questionnaire; PVAQ, Pain and vigilance awareness questionnaire; R, range; SD, standard deviation; $\bar{x}$ , mean.*

### 3.2. Rating of single heat pulses

The number of reported double sensations increased with increasing temperature. Five participants were not able to feel a double sensation at 49°C while all participants felt at least one double sensation at 55°C. On average, when calculating the ratio of fiber specific descriptors across all temperatures to the total amount of descriptors, a greater number of A-δ fiber specific pain descriptors were assigned to the first pain (69.72%), while this pattern was reversed for C-fiber specific descriptors attributed to the second pain (82.89%). An overview of perceived double sensations and pain quality descriptors can be seen in **Appendices 3 - 5.** Reaction times differed significantly between temperatures of 49°C, 51°C, 53°C and 55°C (F (3, 200.6) = 7.02, p < .001, η^2^p = 0.1). Bonferroni-corrected post hoc comparisons showed significantly shorter reaction times between 49°C and 53°C (p < 0.01), 49°C and 55°C (p < 0.01), 51°C and 53°C (p = 0.02) as well as 51°C and 55°C (p = 0.03).

To summarize, the results show that participants were able to clearly distinguish between first and second pain. This differentiation, also regarding A-δ and C-fiber characteristics, could be confirmed by pain quality assessment, whereby the reaction time decreased with increasing heat.

### 3.3. Offset Analgesia

Dependent t-tests showed a significant difference between OT and CT in the T3 interval, indicating an OA response (t (31) = 4.20, p < 0.001, d_z_ = 0.86). Participants reported significantly higher eVAS scores for CT in T3 (x̄ = 25.2, SD = 15.4) compared to OT (x̄ = 12.7, SD = 14.6). An overview of the OA reaction is shown in **Figure 2**.

**Fig 2.**
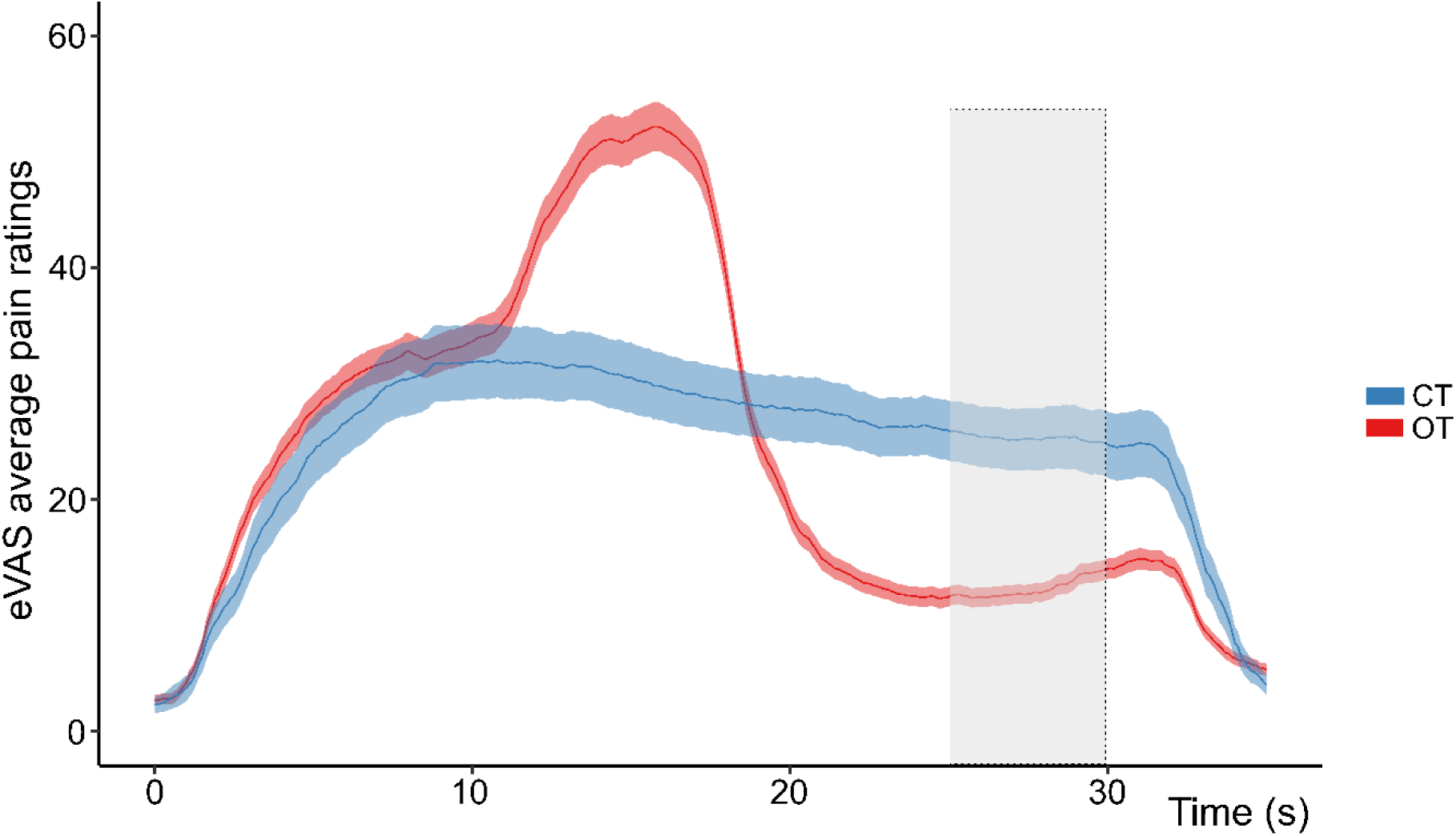
Averaged pain ratings collected during offset and constant trials. Means and standard errors of the mean (SEM) for pain response obtained by an electronic Visual Analogue Scale (eVAS). Offset trial (OT) consisted of an intensity of 46°C for 10 seconds (T1), followed by 5 seconds of 47,5 °C (T2) and ended with a temperature of 46°C for 15 seconds (T3). Constant trials (CT) consisted of an intensity of 46°C for 30 seconds.

### 3.4. Offset analgesia including heat pulses

The majority of participants successfully perceived first and second pain following a heat pulse, with 97% (n=31) demonstrating this perception after two seconds during the T3 interval and 91% (n=29) reporting a perception after 15 seconds during the CTs. Similarly, this was also shown for the OTs (after 2 and after 15 seconds, respectively: n=26, 82%, n=28, 88%). The amount of perceived double sensations for the heat pulse was not significantly different between CT and OT in T3 (15 seconds: W=111.5, p=0.23; 2 seconds: W=84.5, p = 0.14). The number of A-delta fiber pain descriptors that could be assigned to the first pain did not differ between pulses presented during OTs (n=32, 53.3%) and during CTs (n=40, 66.7%) at two seconds (W=90, p=0.064), nor between OTs (n=41, 68.3%) and CTs (n=46, 76.7%) at 15 seconds (W=45.5, p=0.26). A visual representation can be seen in **Figure 3**, for absolute and relative frequencies please see **Appendix 6**.

**Fig 3.**
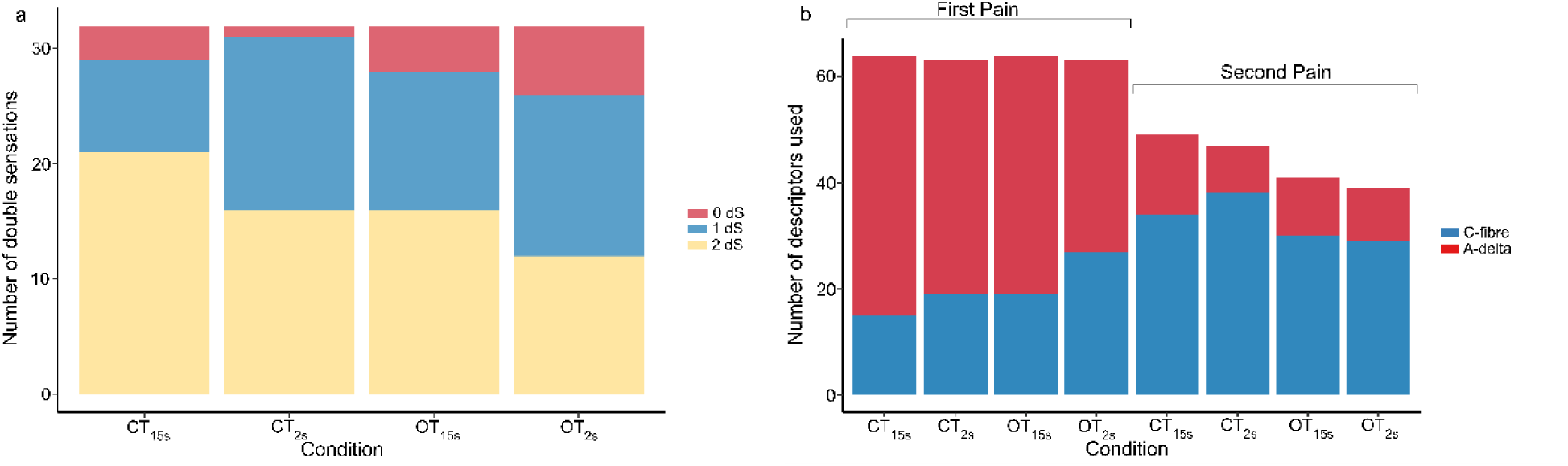
Characteristics of first and second pain during Offset Analgesia. **a)** Number of felt double sensations during offset analgesia in the four conditions. Each condition was tested twice, resulting in a maximum of two felt double sensations per condition. **b)** Number of A-delta or C-fiber descriptors used to describe either the first or second pain. Conditions were either a constant or offset trial with a single heat pulse after two seconds (CT_2s_ / OT_2s_) or after fifteen seconds (CT_15s_ / OT_2s_). The order in which the conditions were tested was randomized.

Regarding the reaction time, a GLM with repeated measures showed a significant interaction of the factors “time” (2 or 15 seconds) and “trial” (OT or CT) (F (1, 211.99) = 4.56, p = 0.03, η^2^p = 0.02). Bonferroni-corrected post hoc comparisons showed significantly higher RTs for OT_2s_ vs CT_2s_ (p < 0.01), OT_15s_ vs CT_2s_ (p < 0.01) as well as CT_15s_ vs CT_2s_ (p = 0.03). Mean reaction times for each trial are shown in **Figure 4**. Regarding the perceived pain intensity, a GLM with repeated measures showed a significant interaction of the factor “time” and “trial” for the first pain, but no significance for the second pain in the Bonferroni-corrected post hoc comparisons. A detailed description of the NRS values and a visual representation can be seen in **Appendix 7**.

**Fig 4.**
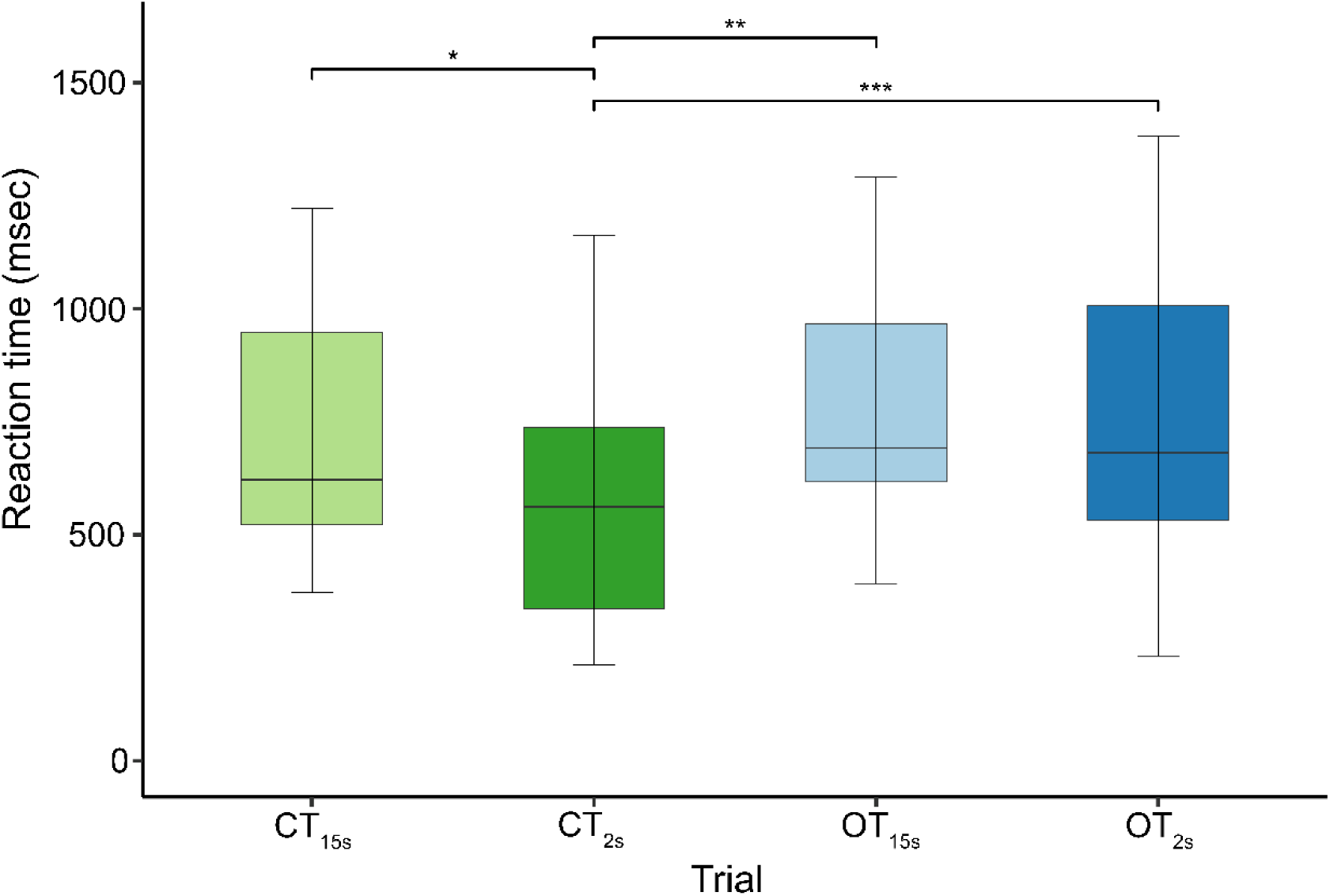
Mean reaction times to heat pulses during offset analgesia. CT with one pulse after 2 seconds (CT_2s_) and one pulse after 15 seconds (CT_15s_). OT with one pulse after 2 seconds (OT_2s_) and one pulse after 15 seconds (OT_15s_). * p<0.05, ** p<0.01, *** p<0.001).

## 4. Discussion

The objective of this study was to examine the potential influence of peripheral fibers on OA represented by first and second pain, which may indicate a role for peripheral pain processing. Contrary to our initial hypothesis, the integration of single heat pulses into an OA paradigm did not reveal significant differences in the analysis of dual sensations and fiber-specific pain descriptors between OTs and CTs across various time intervals. Although response times to heat stimuli were partially prolonged within the OA paradigm, pointing at an involvement of C-fibers rather than A-δ fibers, these findings suggest that peripheral fiber-specific influences exert only a marginal effect on OA processing. Consequently, it is of critical importance to consider potential effects on central mechanisms.

The observed pain responses to brief heat impulses in this study demonstrate that it is feasible to successfully discriminate between first and second pain, as well as qualitatively describe the pain sensation using A-fiber or C-fiber specific descriptors (Beissner et al., 2010; Price et al., 1977; Price, Donald D.; Dubner, 1977). Implementing these concepts into a tonic heat paradigm was successful and showed similar results regarding pain quality ratings and success of discrimination regarding first and second pain. Interestingly, transient change of stimulus intensity - as it happens in the OT - did not influence these results. This is in contrast to earlier research, suggesting an involvement of specific A-δ afferents in OA and does not support the hypothesis of peripheral mechanisms mediating OA (Asplund et al., 2021; Luebke, Von Selle, et al., 2024; Naugle et al., 2013; Szikszay et al., 2022). Indeed, our á priori hypothesis was that the differentiation between first and second pain would be negatively affected by OT. We assumed that OA leads to a temporary A-fiber specific nociceptor fatigue which distorts the sensory experience of first and second pain. Peripheral fatigue of nociceptors in the context of OA has been discussed in previous research as peripheral fatigue has been shown to occur after repeated thermal stimulation of a fixed area in humans (Greffrath et al., 2007) as well as in vitro (Schwarz et al., 2000) and in vivo (Peng et al., 2003) in animal models. The trial specific increases in stimulation intensity, as well as the smaller amount of A-δ nociceptors distributions in the skin, compared to C-fibers (Heppelmann et al., 1988; Ochoa & Mair, 1969), could have influenced the nociceptive response. However, the observed characteristics of first and second pain in this study do not underline a major role of A-fibers in mediating OA. This is supported by a recent study from our lab, in which OA could be induced in participants despite them having a non-ischemic peripheral A-δ nerve block (Luebke, Lopes, et al., 2024). Research that has been focusing on peripheral mechanisms mediating OA, has been mostly based on the theory of glabrous skin not having any A-fiber mechano-heat Type II (AMH-II) nociceptors compared to non-glabrous skin as this has been shown in primates (Campbell & LaMotte, 1983; Meyer et al., 1991; Treede et al., 1995). However, this hypothesis has to be viewed with caution as a similar psychophysical response to heat pain has been shown when stimulating glabrous vs. non-glabrous skin using non-contact heat stimulation (Iannetti et al., 2006) as well as showing OA when stimulating the glabrous skin of the oral mucosa using a contact heat stimulation (Szikszay et al., 2022). These contradicting results may be explained by the difference in epidermal thickness when comparing the glabrous skin of the palm and the non-glabrous skin of the forearm (that is usually used for testing the OA paradigm) instead of the hypothesis of missing AMH-II nociceptors in glabrous skin.

Additional approaches to explain the phenomenon of OA were focusing on the involvement of central nervous system in mediating OA. Several neural imaging studies show the involvement of supraspinal and spinal structures, such as cortical (Alter et al., 2022; Derbyshire & Osborn, 2009; Nahman-Averbuch et al., 2014; Yelle et al., 2009) and subcortical (Derbyshire & Osborn, 2009; Horing et al., 2019; Yelle et al., 2009) structures, cerebellum (Ruscheweyh et al., 2014), brainstem (Derbyshire & Osborn, 2009) and the spinal level (Ligato et al., 2018; Petre et al., 2017; Sprenger et al., 2018). Surprisingly, research trying to confirm these results by using centrally acting drugs such as opioids (Martucci et al., 2012; Niesters et al., 2011, 2014; Olesen et al., 2018; Suzan et al., 2015) opioid-antagonists (Martucci et al., 2012), a serotonin-noradrenalin reuptake inhibitor (Olesen et al., 2018) and an N-Methyl-D-Aspartate receptor antagonist (Bannister et al., 2015; Suzan et al., 2015) were not able to influence the OA response in healthy participants or participants with chronic pain in a significant manner – displaying the robustness of the mechanisms driving OA. Given the variability in the results of these mechanistic approaches, the precise contribution of the identified structures to this phenomenon remains unclear, and further investigation is required to better understand how OA is modulated.

Response time to heat pulses added to an OA paradigm was influenced by trial specific changes of stimulation intensity and duration; however, it remains unclear whether these changes are attributable to OA-specific peripheral mechanisms or other factors, such as the level of nociceptive input. The level of nociceptive input in this context would be determined either by the stimulus intensity, as observed with single heat pulses, or by the duration of the applied stimulation. To be precise, participants exhibited longer RTs in OT compared to CT at both two and fifteen seconds after reaching T3. Given that OA-specific effects typically persist only for a few seconds (Yelle et al., 2008), it is more likely that mechanisms unrelated to OA underlie these changes. Additionally, within CTs, RT increased for longer CTs (terminated 15 seconds after T3) compared to shorter CTs (terminated 2 seconds after T3), further supporting the hypothesis that RT alterations were not driven by OA-specific peripheral mechanisms but rather by factors such as the overall level of nociceptive input. This finding aligns with previous research demonstrating that continuous painful stimulation leads to prolonged RT (Attridge et al., 2016; van Laarhoven et al., 2017).

An explanation for the difference in pain quality measurements and RT may be that the outcome measurements involve distinct mechanisms of processing. Differentiating and interpreting a double pain sensation may rely more on conscious processing, enabling discrimination of sensory inputs regardless of afferent information speed. The perception and processing of pain is much more complex, with contributions from multiple levels of the neuroaxis (Coghill et al., 1999; Fenton et al., 2015; Koyama et al., 2005; Sgourdou, 2022; Yang & Chang, 2019). In contrast, reacting to a painful stimulus has been shown to depend heavily on afferent transduction and maybe does not depend on a full awareness of the stimulus (Castellote & Valls-Solé, 2019; Henderson & Dittrich, 1998; Yarnitsky & Ochoa, 1991). Several factors may have influenced RT changes, including alterations in afferent and efferent transmission, central processing affected by pain or OA mechanisms, and limitations in attention or cognitive resources - specifically, characteristics of the network or its modulators(Lakhani et al., 2012). These factors may interact with pain-specific central processing, as brain regions, including the prefrontal cortex, have been shown to be involved in processing tonic heat stimuli and the modulation attention (Alba et al., 2022; Nahman-Averbuch et al., 2014; Ong et al., 2019; Schulz et al., 2015).

### 4.1. Limitation and future direction

Whether specific peripheral nerve fibers play a role in OA cannot be conclusively clarified with this study, since this would require a thorough investigation of electroneurographic test procedures to identify specific nerve fibers. Nonetheless, we chose a rigorous approach by investigating several qualitative and quantitative aspects of pain, as well as observing outcome measures separate to pain (RT), to depict the effect of OA on peripheral nerve fibers as thoroughly as possible. When looking at average eVAS pain ratings it is apparent that our stimulation intensities may not have been intense enough to elicit a strong offset effect, especially in contrast to the CT which shows average eVAS ratings below 40/100. This could’ve influenced the effect of the OA paradigm on our quantities of interest, and it may have been more robust to calibrate stimulus intensity to each individual. This question is a common methodological issue (Adamczyk et al., 2022) and needs to be taken into account, however an influence on our results is highly unlikely, since we were able to show a significant offset effect. Furthermore, the same stimulus intensity may be better suited to produce equal afferent activation patterns as discussed and shown in prior research (Adamczyk et al., 2022; Coghill et al., 2003; Sprenger et al., 2015). Additionally, previous studies have been using this combined approach to disentangle peripheral from central response times (Lakhani et al., 2012; Ploner et al., 2006).

In summary, this study demonstrates that the integration of single heat pulses into an OA paradigm did not reveal significant differences in the analysis of double sensations or fiber-specific pain descriptors between an offset analgesia-paradigm and a control condition across various time intervals. These findings suggest that central mechanisms may play a more prominent role in the modulation of OA, with peripheral fiber-specific effects exerting limited influence. Future research should focus on investigating OA-specific central mechanisms to further elucidate the neurophysiological processes underlying offset analgesia and its impact on pain perception.

## 5. Statements and Declarations

The authors declare that they have no conflict of interest. This study is supported by the Deutsche Forschungsgemeinschaft (DFG, German Research Foundation)—493000854.

