## Appendix (supplementary materials) for "Perception of first and second pain during offset analgesia"

**S 1: Presentation and distinction of first and second pain**

Before the start of data collection, each participant was instructed to follow a Microsoft PowerPoint presentation (Microsoft Corp., Remond, WA, USA) that described the study procedure, the phenomenon of first and second pain, the testing methods and the eVAS. First and second pain were explained with written and visual information including a chart with associated pain descriptors.

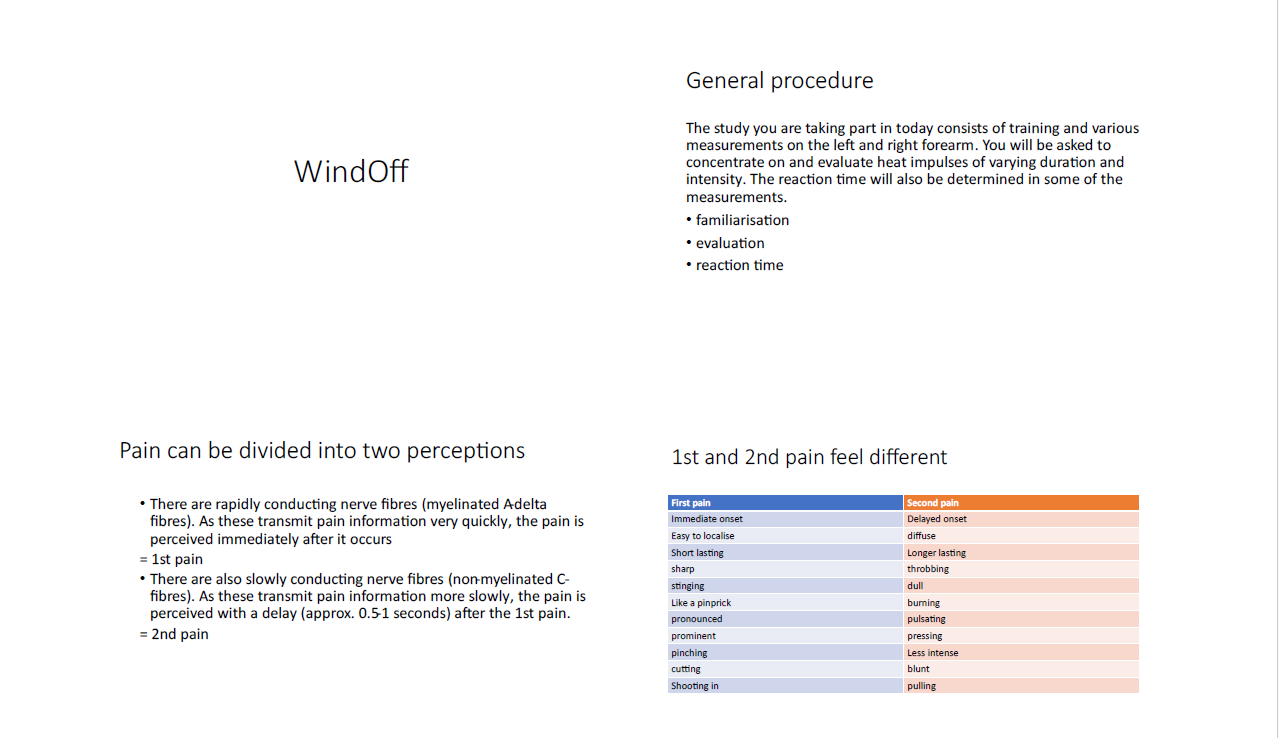

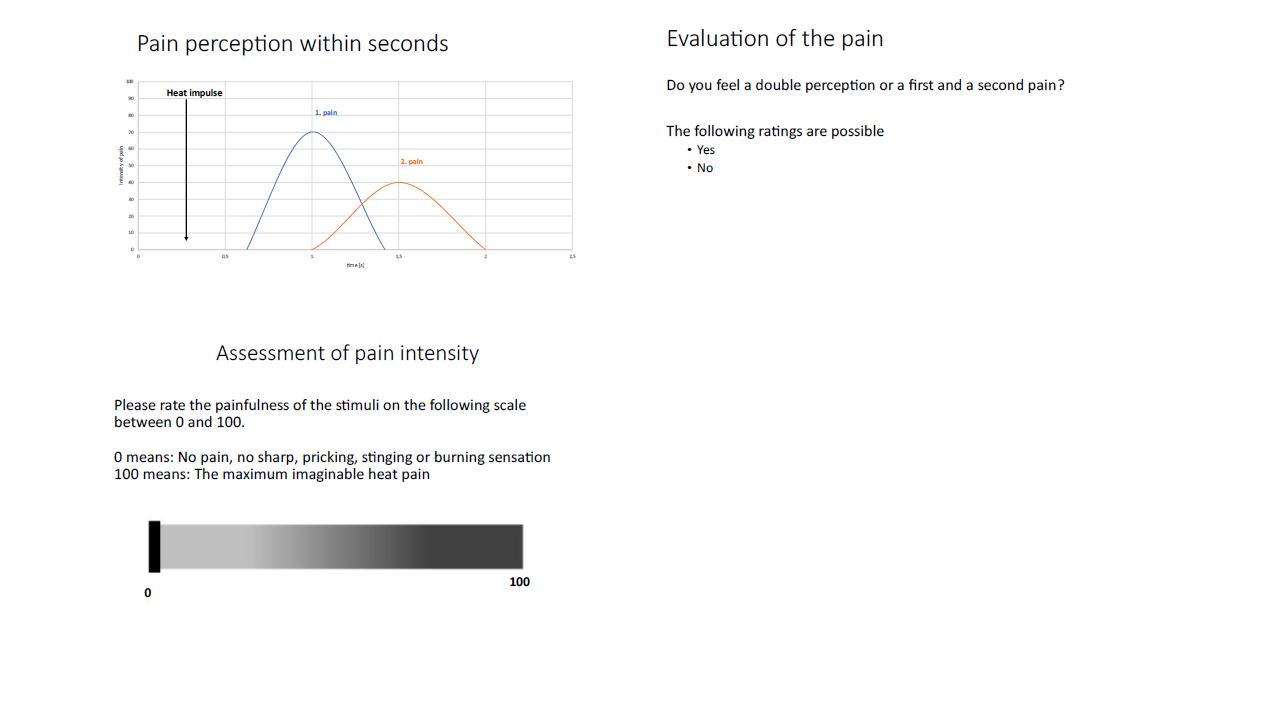

**S 2: Pain descriptors for first and second pain**

A literature search was conducted to identify pain descriptors in PubMed dealing with first and second pain triggered by heat stimuli: (first pain) OR (second pain) AND heat). These pain descriptors were systematically extracted and presented to participants in a) a sorted manner during preparation and b) a randomized manner during the consecutive study.

**2a: sorted**

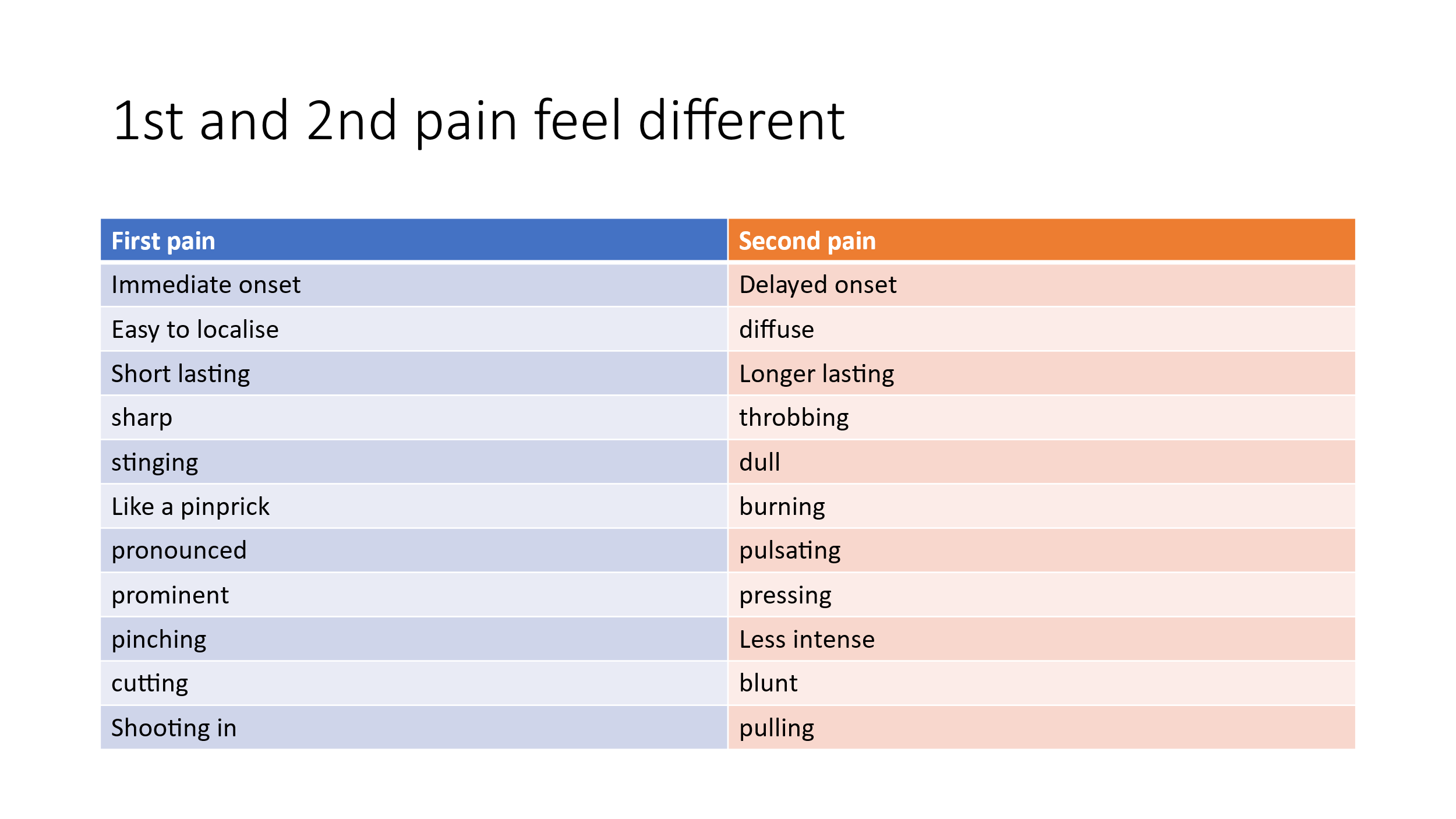

***2b: randomized***

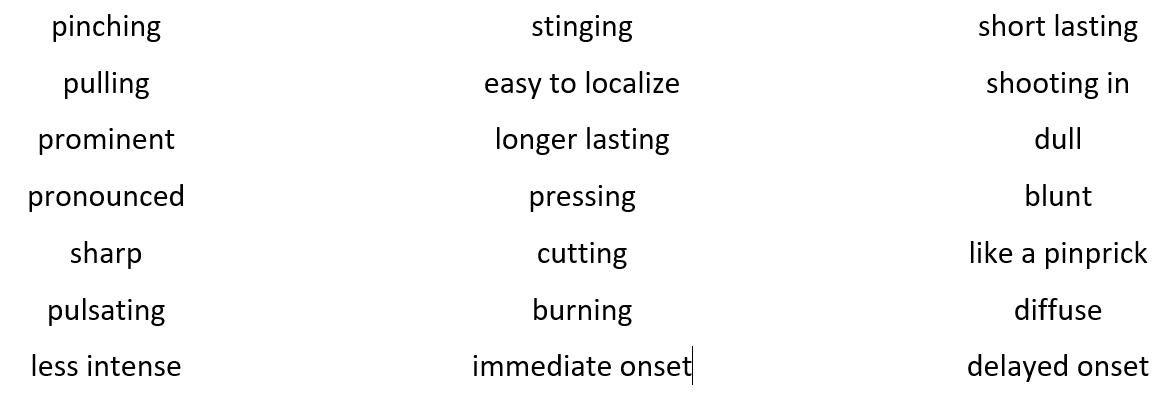

**S 3: Rating of single heat pulses: Perceived double sensations**

| **Temperature [°C]** | **No double**  **sensation** | **One double sensation** | **Two double sensations** |
| --- | --- | --- | --- |
| **49** | 5 (16%) | 16 (50%) | 11 (34%) |
| **51** | 4 (13%) | 8 (25%) | 20 (63%) |
| **53** | 2 (6%) | 7 (22%) | 23 (72%) |
| **55** | 0 (0%) | 8 (25%) | 24 (75%) |

The values are indicated in absolute and relative frequency. Each temperature (49°C, 51°C, 53°C, 55°C) has been tested twice, therefore a maximum of two double sensations per temperature was possible.

**S 4: A-fiber and C-fiber pain descriptors for single heat pulses shown in absolute and relative frequencies.**

|  | | **First pain** | | | | **Second pain** | | | |
| --- | --- | --- | --- | --- | --- | --- | --- | --- | --- |
| **Pain quality** | | 49°C | 51°C | 53°C | 55°C | 49°C | 51°C | 53°C | 55°C |
| **A-δ-fibre** | Stinging | 14 | 12 | 28 | 18 | 1 | 2 | 3 | 2 |
|  | sharp | 5 | 5 | 5 | 5 | 0 | 1 | 0 | 1 |
|  | Like a pinprick | 1 | 7 | 3 | 5 | 2 | 1 | 0 | 1 |
|  | Easy to localise | 5 | 3 | 3 | 3 | 1 | 2 | 0 | 0 |
|  | cutting | 2 | 2 | 4 | 3 | 1 | 0 | 0 | 1 |
|  | pinching | 4 | 3 | 0 | 1 | 1 | 0 | 1 | 0 |
|  | Short lasting | 4 | 2 | 4 | 1 | 2 | 2 | 1 | 1 |
|  | Shooting in | 1 | 2 | 2 | 9 | 0 | 0 | 0 | 1 |
|  | prominent | 1 | 2 | 0 | 0 | 0 | 0 | 0 | 1 |
|  | pronounced | 1 | 3 | 0 | 1 | 0 | 0 | 0 | 0 |
|  | Immediate onset | 1 | 0 | 0 | 0 | 0 | 1 | 1 | 1 |
|  | **Sum of Aδ-fibre descriptors** | **61% (39)** | **65% (41)** | **78% (49)** | **75% (46)** | **22%**  **(8)** | **19%**  **(9)** | **12%**  **(6)** | **17%**  **(9)** |
| **C-fibre** | dull | 0 | 1 | 1 | 0 | 4 | 6 | 5 | 8 |
|  | Less intense | 6 | 6 | 3 | 4 | 4 | 5 | 6 | 6 |
|  | diffuse | 2 | 3 | 0 | 0 | 7 | 8 | 8 | 4 |
|  | pulling | 5 | 1 | 2 | 1 | 2 | 4 | 6 | 2 |
|  | Delayed onset | 3 | 0 | 1 | 1 | 2 | 8 | 4 | 6 |
|  | blunt | 0 | 1 | 1 | 0 | 1 | 1 | 2 | 3 |
|  | burning | 5 | 9 | 6 | 9 | 5 | 4 | 12 | 10 |
|  | throbbing | 1 | 1 | 0 | 0 | 1 | 0 | 0 | 0 |
|  | pressing | 1 | 0 | 0 | 0 | 1 | 0 | 0 | 0 |
|  | pulsating | 0 | 0 | 0 | 0 | 0 | 0 | 0 | 0 |
|  | Longer lasting | 2 | 0 | 0 | 0 | 1 | 2 | 3 | 4 |
|  | **Sum of C-fibre descriptors** | **35% (25)** | **35% (22)** | **22% (14)** | **25% (15)** | **78% (28)** | **81% (38)** | **88% (46)** | **83% (43)** |

Relative frequencies were determined by dividing the number of descriptors felt per temperature and fiber type by the total number of descriptors per temperature.

**S 5 A and B**

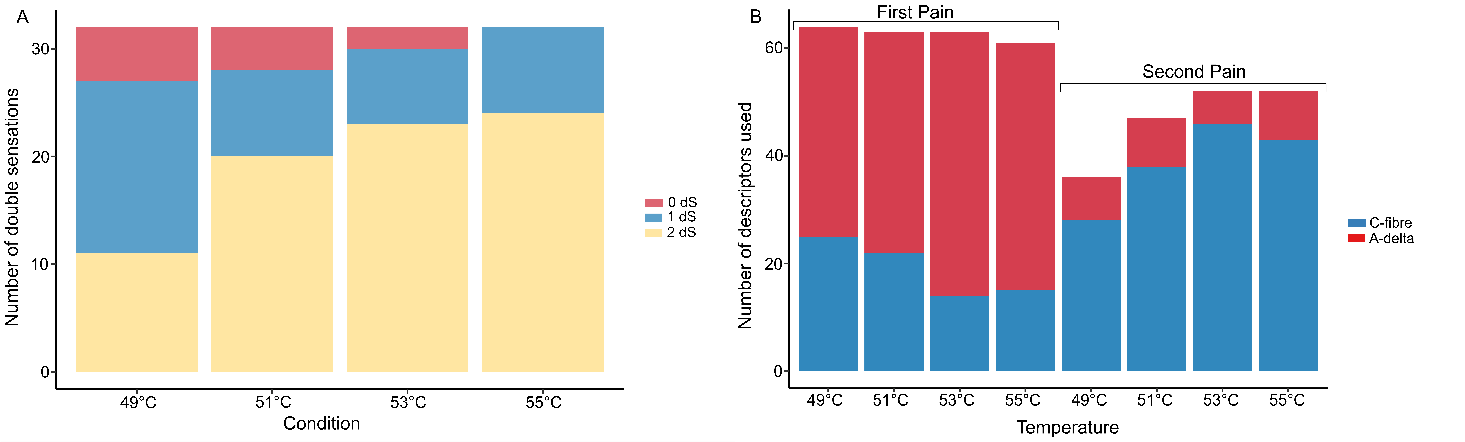

**Fig. 5. Qualitative properties of first and second pain during single heat pulses.** **A:** Number of felt double sensations from single heat pulses in the four temperatures. Each condition was tested twice, resulting in a maximum of two felt double sensations per temperature. **B:** Number of A-delta or C-fiber descriptors used to describe either the first or second pain. Temperatures tested were 49°C, 51°C, 53°C and 55°C. The applied order of single heat pulses was randomized.

**S 6: Perceived double sensations for offset analgesia including heat pulses**

A total of four conditions were tested, consisting of either an offset trial or constant trial with including a heat pulse after either two seconds (CT_2s_ / OT_2s_) or after fifteen seconds (CT_15s_ / OT_15s_).

***6a:* Double sensations**

| **Temperature [°C]** | **No double**  **sensation** | **One double**  **sensation** | **Two double sensations** |
| --- | --- | --- | --- |
| **CT_15s_** | 3 (9%) | 8 (25%) | 21 (66%) |
| **CT_2s_** | 1 (3%) | 15 (47%) | 16 (50%) |
| **OT_15s_** | 4 (13%) | 12 (38%) | 16 (50%) |
| **OT_2s_** | 6 (19%) | 14 (44%) | 12 (38%) |

Relative frequencies were determined by dividing the number of sensations felt per condition by the total number of double sensations per condition.

***6b:* A-fiber and C-fiber pain descriptors for offset analgesia including heat pulses shown in absolute and relative frequencies.**

|  | | **First pain** | | | | **Second pain** | | | |
| --- | --- | --- | --- | --- | --- | --- | --- | --- | --- |
| **Pain quality** | | CT_15s_ | CT_2s_ | OT_15s_ | OT_2s_ | CT_15s_ | CT_2s_ | OT_15s_ | OT_2s_ |
| **A-δ-fibre** | Stinging | 17 | 19 | 16 | 15 | 3 | 3 | 2 | 5 |
|  | sharp | 4 | 5 | 3 | 6 | 3 | 1 | 0 | 0 |
|  | Like a pinprick | 7 | 2 | 3 | 0 | 0 | 1 | 1 | 0 |
|  | Easy to localise | 2 | 2 | 4 | 4 | 1 | 0 | 0 | 1 |
|  | cutting | 3 | 2 | 4 | 3 | 2 | 1 | 1 | 1 |
|  | pinching | 1 | 1 | 3 | 2 | 2 | 1 | 2 | 1 |
|  | Short lasting | 2 | 0 | 2 | 3 | 0 | 0 | 1 | 1 |
|  | Shooting in | 7 | 10 | 8 | 2 | 3 | 1 | 1 | 0 |
|  | prominent | 2 | 2 | 1 | 0 | 1 | 0 | 0 | 0 |
|  | pronounced | 2 | 1 | 1 | 0 | 0 | 1 | 2 | 1 |
|  | Immediate onset | 2 | 0 | 0 | 1 | 0 | 0 | 1 | 0 |
|  | **Sum of Aδ-fibre descriptors** | **77% (49)** | **70% (44)** | **70% (45)** | **57% (36)** | **31% (15)** | **19%**  **(9)** | **27% (11)** | **26% (10)** |
| **C-fibre** | dull | 0 | 0 | 0 | 2 | 5 | 1 | 9 | 5 |
|  | Less intense | 1 | 6 | 1 | 3 | 4 | 4 | 3 | 5 |
|  | diffuse | 1 | 1 | 2 | 2 | 6 | 7 | 7 | 5 |
|  | pulling | 1 | 1 | 2 | 4 | 8 | 6 | 3 | 4 |
|  | Delayed onset | 0 | 0 | 1 | 1 | 2 | 3 | 1 | 3 |
|  | blunt | 0 | 2 | 2 | 1 | 0 | 2 | 0 | 1 |
|  | burning | 12 | 6 | 10 | 10 | 7 | 13 | 4 | 5 |
|  | throbbing | 0 | 0 | 0 | 1 | 1 | 1 | 1 | 1 |
|  | pressing | 0 | 0 | 0 | 0 | 0 | 0 | 2 | 0 |
|  | pulsating | 0 | 0 | 0 | 0 | 0 | 0 | 0 | 0 |
|  | Longer lasting | 0 | 3 | 1 | 3 | 1 | 1 | 0 | 0 |
|  | **Sum of C-fibre descriptors** | **23% (15)** | **30% (19)** | **30% (19)** | **43% (27)** | **69% (34)** | **81% (38)** | **73% (30)** | **74% (29)** |

Relative frequencies were determined by dividing the number of descriptors felt per temperature and fibre type by the total number of descriptors per temperature.

**S 7: Perceived intensity of first and second pain in the OA paradigm**

To investigate perceived intensity of first and second pain the mean pain intensity for each condition (CT_15s,_ CT_2s,_ OT_15s,_ OT_2s_, two trials per condition) were calculated. Furthermore, a generalized linear model (GLM) (2 x 2 design), was used. If there were significant findings, Bonferroni-corrected t-tests were performed. The level of significance was set at p < 0.05.

The GLM with repeated measures showed a significant interaction of the factor “time” (2 or 15 seconds) and the factor “trial” (OT or CT) for first pain (F (1, 93) = 10.90, p < 0.01, η^2^p = 0.10), but not for the second pain (F (1, 79.74) = 6.39, p = 0.01, η^2^p = 0.07). For the first pain, Bonferroni corrected group comparisons revealed a significant lower NRS rating for OT_2s_ vs. CT_2s_ (p < 0.001) and a significant higher NRS rating for CT_15s_ vs. OT_2s_ (p < 0.01) and OT_15s_ vs. OT_2s_ (p < 0.01). The results for the second pain showed no significance after Bonferroni corrected post-hoc testing.

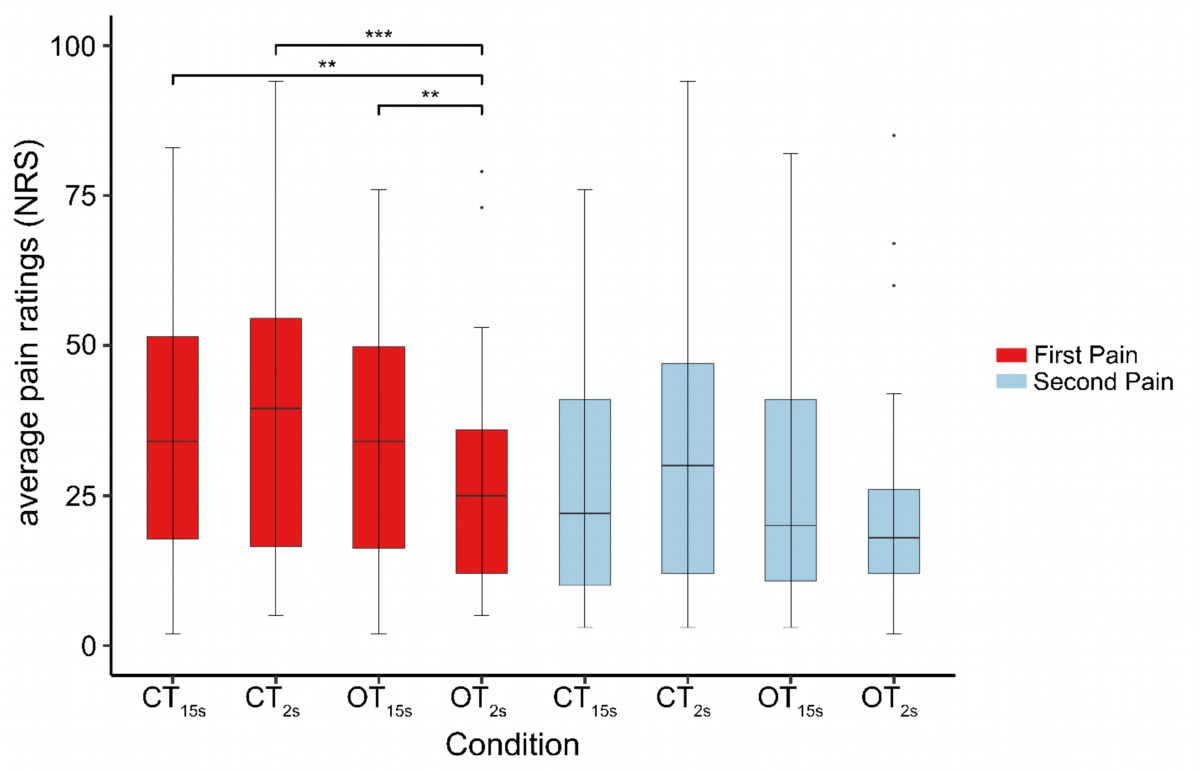

**Fig. 6. Perceived intensity of first and second pain in the offset analgesia paradigm.** CT with one pulse after 2 seconds (CT_2s_) and one pulse after 15 seconds (CT_15s_). OT with one pulse after 2 seconds (OT_2s_) and one pulse after 15 seconds (OT_15s_). Pain ratings were collected on a numerical rating scale (NRS) from 0 – 100. Participants evaluated their first pain (red) and second pain (blue) in each trial. * p<0.05, ** p<0.01, *** p<0.001)
